# A radial pillar device (RAPID) for continuous and high-throughput separation of multi-sized particles

**DOI:** 10.1101/197046

**Authors:** Ninad Mehendale, Oshin Sharma, Claudy D’Costa, Debjani Paul

## Abstract

Pillar-based microfluidic sorting devices are preferred for isolation of rare cells due to their simple designs and passive operation. Dead-end pillar filters can efficiently capture large rare cells, such as, circulating tumor cells (CTCs), nucleated red blood cells (NRBCs), etc., but they get clogged easily. Cross flow filters are preferred for smaller rare particles (e.g. separating bacteria from blood), but they need additional buffer inlets and a large device footprint for efficient operation. We have designed a new microparticle separation device i.e. <u>Ra</u>dial <u>Pi</u>llar <u>D</u>evice (RAPID) that combines the advantages of dead-end and cross flow filters. RAPID can simultaneously isolate both large and small rare particles from a mixed population, while functioning for several hours without clogging. We have achieved simultaneous separation of 10*µ*m and 2*µ*m polystyrene particles from a mixture of 2 *µ*m, 7 *µ*m and 10 *µ*m beads. RAPID achieved average separation purity and recovery in excess of ⟂ 90%. The throughput of our device (⟂ 3ml/min) is 10 and 100 times higher compared to cross flow and dead-end filters respectively, thereby justifying the name RAPID.

## 1 Introduction

One of the simplest ways to isolate microparticles from a heterogeneous mixture with minimal pre-processing is to separate them according to their size. Various pillar-based microfluidic separation strategies have been reported in the literature, such as, dead-end filters [1], cross flow filters[2], deterministic lateral displacement (DLD) device [3], OncoBean [4], ratchetbased separation [5], etc. Dead-end pillar devices (fig. 1B) are designed with pillar gaps that are slightly smaller than the size of the particles to be trapped. This allows smaller particles to easily pass through the pillar network, while the larger particles are trapped. The trapped large particles then tend to stack, eventually stopping the device operation within a short time. [6, 7, 8, 9].

**Figure 1:**
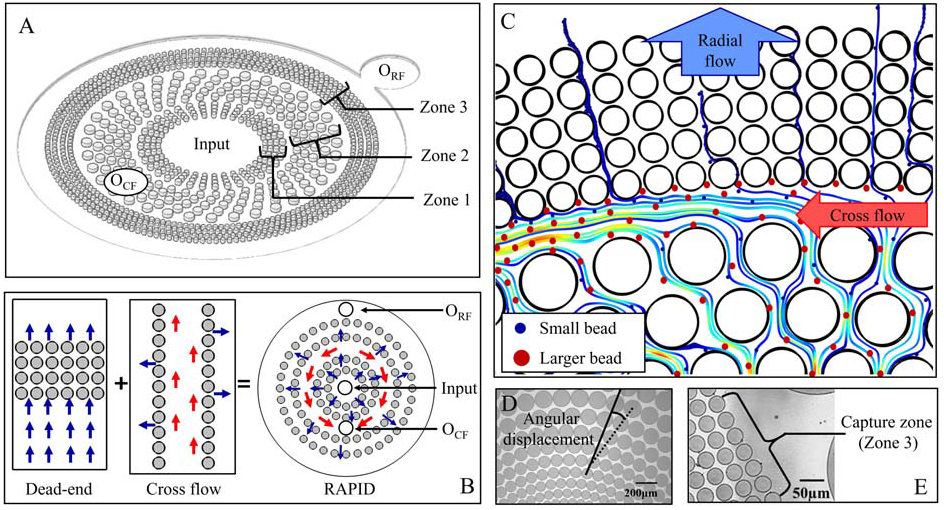
Principle of operation of RAPID. (A) Schematic of RAPID. Device has a central inlet which is also used to collect large particles after the experiment. RAPID has a cross flow outlet (O_*CF*_) for intermediate-sized waste particles and a radial flow outlet (O_*RF*_) for small particles. The pillars are arranged in concentric circles in three zones, with the pillar gap varying across each successive zone. Zone 1 is a pre-filter that traps large particles. The cross flow is set up in zone 2. Zone 3 allows the smaller particles to go towards O_*RF*_, while blocking the intermediate-sized particles. (B) Fluid flows into the pillars in a dead-end filter. In a cross flow filter, large particles follow the cross flow (red arrows), while the smaller particles follow the side flow into the pillars (blue arrows). RAPID starts as a radial flow device (blue arrows) similar to a dead-end filter, and switches to a cross flow (red arrow) operation at the onset of clogging. This is achieved by displacing the successive rows of pillars by a pre-determined angle. (C) Schematic diagram showing two kinds of flows in RAPID at the intersection of zones 2 and 3. Smaller beads (blue) follow the radial path. Intermediate-sized beads (red) cannot flow into zone 3 and follows the tangential cross flow path. (D) Microscope image of the device showing the angular displacement (*θ*) between successive rows in zone 2. (E) Microscope image of the pillars in zone 3.

To address the clogging problem, some groups [10, 5] shaped the pillars like ratchets and reversed the flow periodically. Other groups integrated micro-pumps [11], performed pneumatic actuation [12, 13, 14] or introduced mechanical vibration using piezoelectric transducers [15]. All these approaches required complex micro-fabrication or the integration of power sources and transducers.

Chen *et al*. [16] designed a cross flow filter (fig. 1B). Here, the pillars are arranged along the length of the channel (i.e. perpendicular to the main flow), leading to passive bifurcation of the flow. Clogging is avoided as the larger particles are carried away by the main flow. As most particles prefer the path of low hydrodynamic resistance [17], many small target particles also remain in the main channel along with the large particles. Hence, cross flow filters are not suitable for isolation of large rare cells [18, 19]. Li *et al*. [20] proposed a pulsatile flow to increase the efficiency of cross flow micro-filtration. Geng and others [21] arranged their cross flow channel in a spiral to reduce the device footprint. As the separated particles have to travel the entire spiral path after being sorted, there was a chance of remixing due to Dean flows. They avoided remixing using a split-level design in a silicon chip but requires complex two-level lithography and silicon etching.

We have designed a microparticle enrichment device (i.e. <u>Ra</u>dial <u>Pi</u>llar <u>D</u>evice or RAPID) (fig. 1) that combines the advantages of dead-end and cross flow filters. RAPID does not need a separate buffer inlet and can function for several hours. Two design features help in continuous operation of this device. First, the pillars are arranged in successive concentric circles in three different zones. Due to the arrangement of pillars along a circle, a higher number of pillars can be accommodated in the first row itself compared to a typical dead-end device with the same footprint. This significantly delays the onset of clogging, as there are multiple parallel paths available for the particles. Second, the pillars in the successive rows are displaced by a pre-determined angle (*θ*) to generate a cross flow towards waste outlet (O_*CF*_). The large rare particles are trapped by the pillars of zone 1, while the intermediate-sized and small particles continue to flow into the device. The intermediate-sized particles are removed by the cross flow, while small particles continue to move radially out of the device. These two features enhance the throughput (3 ml/min) and lead to continuous device operation, justifying the name RAPID.

## 2 Materials and methods

### 2.1 Equipment and chemicals

SU-8 2010 and its developer were obtained from MicroChem Corporation (West-borough, USA). Sylgard-184 polydimethylsiloxane (PDMS) was purchased from Dow Corning Corporation (Michigan, USA). 40mm x 24 mm glass cover slips (No. 1) were obtained from Blue Star, Mumbai, India. Microfluidic connectors (barb-to-barb WW30626-48 and luer-to-barb WW-30800-06) were bought from Cole-Parmer (Mumbai, India). Tygon tubing of 1.5 mm diameter (formulation 2375) was used to connect the chip to the syringes. 2*µ*m amine-modified fluorescent polystyrene microparticles, (Sigma L9529), 7*µ*m (Sigma 78462) and 10 *µ*m (Sigma 61946) polystyrene microparticles and Tween-20 surfactant were obtained from Sigma Aldrich (Mumbai, India). 10 ml plastic syringes were obtained from Becton-Dickinson (Mumbai, India). Unless stated, all chemicals were used without any further purification.

UV exposure of photoresist was performed using a MJB4 mask aligner from Karl-Suss. The height of the pillars was measured using an Ambios-XP2 profilometer. Polydimethylsiloxane (PDMS) devices were bonded to glass cover slip using a Harrick plasma cleaner (PDC 32G). A syringe pump (model 111, Cole-Parmer) was used to control the flow. Microparticles were counted using a hemocytometer (Rohem, Nashik, India). Images and videos were acquired using a Nikon Eclipse Ti inverted fluorescence microscope, fitted with 20x and 40X (1.3 NA) objectives and FITC filter.

### 2.2 Device design

The schematic diagrams in fig.1 A and B show the design concept of RAPID. As shown in the figure, the device has a single central inlet, a cross flow outlet and a radial flow outlet. The radial and cross flow outlets are positioned on diametrically opposite sides of the central inlet. The bead sizes were chosen to be 10 *µ*m, 7 *µ*m and 2 *µ*m in this work to demonstrate separation in RAPID. The pillars in this device are arranged in three distinct zones. The pillar gaps (and consequently the pillar diameters) in the three zones were set according to the particle sizes to be separated. There are 9 rows of pillars in the innermost zone 1 (i.e. pre-filter zone) with a gap of 8*µ*m between the pillars. The first row of pillars has a diameter of 50 *µ*m, and the diameter is increased by approximately 10% for each successive row to keep the pillar gap constant throughout this zone. The pillar gap in zone 1 was chosen such that it can trap large rare beads and bead aggregates.

The outermost zone (zone 3) has pillar gaps of 3 *µ*m and pillar diameters starting from 25 *µ*m. There are 39 rows of pillars, with the pillar diameter increasing by 3% for each successive row. The rationale behind choosing such a large number of rows was to ensure that the intermediate-sized beads do not escape from the radial outlet. This zone can enrich and separate particles smaller than 3 *µ*m.

The pillars in zone 2 have a gap of 12 *µ*m, with the diameters increasing radially from 80 to 100 *µ*m over 11 rows. The pillar gap was large to allow all particles coming out of zone 1 to pass through. Each successive row of pillars in zone 2 is given an angular displacement of 5^º^ (fig.1 D) to facilitate the cross flow in this zone. This zone also houses the cross flow (waste) outlet to remove intermediate-sized particles from the device. The enriched sample consisting of small particles is collected from the radial outlet (O_*RF*_). Figures 1D and 1E show microscope images of pillars in zones 2 and 3 respectively.

### 2.3 Optimization of device design by COMSOL simulation

The flow of water in three different design iterations of RAPID was simulated using the COMSOL Multiphysics software (version 5.2) to identify the one with the highest cross flow. We used the built-in microfluidics module (single phase laminar flow) with "no slip" boundary conditions. A flow rate of 1 ml/min at the inlet was chosen. Since the actual device had more than 9000 pillars, and was largely flat with a high width-to-height ratio, we made certain simplifications during simulation. The number of rows in zones 1, 2 and 3 was reduced to 4, 5 and 4 respectively to qualitatively capture the essential flow behavior. Further, the device was simulated in 2D instead of 3D. These simplifications were made to work within the limited computer memory. For the purpose of simulation, the entire peripheral area after zone 3 was treated as the outlet for capturing the small particles.

### 2.4 Device fabrication

The mask was designed in CleWin 4 and prepared on an iron oxide mask plate using a laser writer. The device was fabricated using standard soft-lithography process in the IIT Bombay Nanofabrication Facility. In brief, 2-inch p-type <111> silicon wafers were cleaned by RCA technique and baked on a hot plate at 120°C for 20 minutes to dehydrate them. SU-2010 was spin coated with the following two-step protocol: (i) a spreading spin of 500 rpm for 15 sec, with 200 rpm/min acceleration, and (ii) a final spin of 2500 rpm for 45 sec, with 200 rpm/min acceleration. Soft bake was performed on a hot plate at 95°C for 3 min, followed by cooling to room temperature for 5 min. UV exposure of the pattern was performed for 9 sec (115 mJ/cm2), followed by post bake at 95°C for 5 min. The pattern was developed in SU-8 developer, rinsed in IPA and dried under nitrogen.

Sylgard 184 parts A and B were mixed well in standard 1:10 ratio, degassed and poured on the silicon master. The PDMS was allowed to cure in an oven at 65°C for 1 h. The devices were cut from the cured mold. The inlet and outlets were punched using a 1.5mm biopsy punch. The devices were then bonded using air plasma for 90 sec. The devices were used without any surface treatment. The pillar height was measured to be 10 *µ*m by profilometry.

### 2.5 Sample preparation

We prepared the sample by mixing polystyrene microparticles of three different sizes (10 *µ*m, 7*µ*m and 2*µ*m). The beads were stored at 4°C and allowed to attain room temperature for 20 min before an experiment. The stock bead samples were sonicated for 5 min and subsequently vortexed for 5 min to disperse any aggregates. To avoid microparticle aggregation, 0.1% Tween-20 was added to the particle mix. The final concentrations of the 10 *µ*m, 7 *µ*m, and 2*µ*m beads were 10^3^/l, 3 x 10^7^/l and 3 x 10^7^/l respectively. It should be noted that the concentration of the smaller beads (7 *µ*m and 2 *µ*m) was chosen to be 30,000 times higher than the concentration of the larger beads to demonstrate the capture of large rare particles in RAPID.

We also demonstrated high concentration separation of small rare particles by separating 1 *µ*m beads from a background of 7*µ*m beads. The final concentrations of the 7*µ*m and 1*µ*m beads in DI water were 2.13 x 10^6^/ml and 2.13 x 10^5^/ml respectively. There were approximately ten 7*µ*m beads for every 1 *µ*m bead in the final sample mixture.

### 2.6 Microparticle sorting experiment

10 ml of the mixed bead sample was loaded into a 10 ml syringe and mounted on the syringe pump. The sample was pumped with flow rates of 100 *µ*l/min, 300 *µ*l/min, 500 *µ*l/min, 1000 *µ*l/min and 3000 *µ*l/min respectively. We continued flowing sample into the device until the entire sample volume of 10 ml was exhausted. We noted the time whenever the sample volume increased by 500 *µ*l in the collection tube attached to an outlet. This was done for both outlets to calculate throughput and recovery rate.

After the experiment, the trapped 10 *µ*m beads at the inlet were recovered by reverse flow of 10 ml DI water simultaneously from both outlets. For each flow rate, three separate runs were performed and the beads were counted. The samples were counted three times on a hemocytometer (using 20*µ*l sample volume) under a microscope fitted with a 40X oil immersion objective and then averaged. The beads were sonicated and vortexed for 5 min each before counting. The larger (10 *µ*m and 7 *µ*m) beads were imaged under the brightfield and the 2 *µ*m beads were imaged using fluorescence (FITC filter). Each square of the hemocytometer was separately imaged during counting. A bead counting algorithm in MATLAB was specifically written by us to count the number of each kind of bead from the acquired images.

## 3 Results and discussion

### 3.1 Fluid flow inside RAPID

We went through several design iterations of RAPID before choosing the final design. Fluid flow in each of these devices was simulated using COMSOL prior to device fabrication. Figure 2 shows the simulation results. For all the panels, the top row shows the magnitude of radial flow velocity (v_*r*_), and the bottom row shows the magnitude and direction of the cross flow velocity (v_*θ*_). The design in panel A has a simple radial arrangement of pillars and no cross flow outlet (O_*CF*_). As expected, the fluid flow is radially symmetric and there is no cross flow anywhere in the device. Compared to a simple dead-end filter, the onset of clogging would be delayed in this radial design due to the presence of a large number of parallel paths. As there is no outlet to remove the particles trapped between zones 2 and 3, the device would eventually stop functioning due to particle stacking.

**Figure 2:**
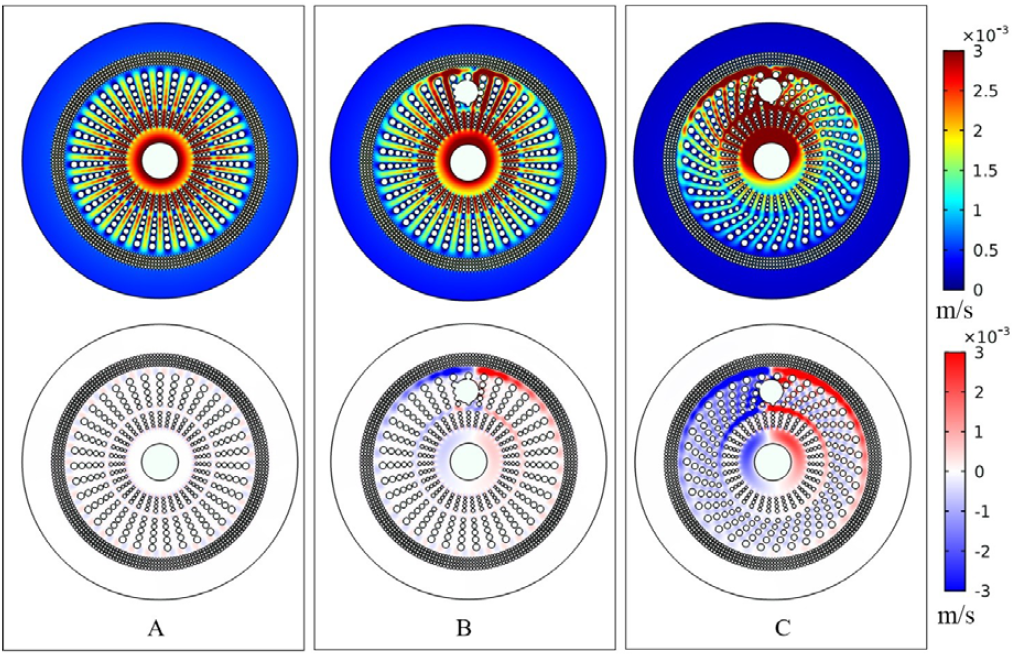
COMSOL simulation results obtained from three design iterations of RAPID. The upper row shows the surface plots of the radial velocity in the device, while the lower row shows the surface plots of the cross flow velocity. For the lower panel, the colors blue and red indicate flow velocity along clockwise and anti-clockwise directions respectively. A white area (except at inlet and outlets) indicates a dead zone, i.e. an area with no net cross flow. (A) The first design had no cross flow outlet (O_*CF*_) and no angular displacement between the successive rows of pillars. As expected, there was strong and symmetric radial flow in the device, with no cross flow. (B) The second design had an additional waste outlet O_*CF*_ in zone 2. Its presence generated a cross flow towards the outlet and increased the radial flow in the device. (C) In the final design, successive rows of pillars in zone 2 were displaced by a pre-determined angle(*θ*). Both radial and cross flows around the outlet were significantly higher compared to the previous design iterations. The cross flow was primarily clockwise following the alignment of the pillars. Design C was chosen for further experiments.

Therefore, in panel B we introduced an outlet (O_*CF*_) in zone 2 to take out the stacked particles. This generated a tangential flow (v_*θ*_) towards the outlet O_*CF*_, in addition to the radial flow. Blue and red indicate clockwise and anticlockwise flows respectively. However, there were still parts of the device (e.g. white areas in the velocity surface plots) with no net cross flow. While stacking would be less here compared to the previous design, some clogged particle layers would still be stacked in these areas.

In the next design iteration (panel C), we introduced a pre-determined angular displacement (*θ*) between the successive rows in zone 2 to give a net directionality to the cross flow and have a non-zero v_*θ*_ component everywhere. This led to a self-circulating cross flow throughout the device with almost zero dead volume. The v_*θ*_ component in this design is primarily clockwise due to the alignment of the pillars. The radial flow was also stronger around the outlet (O_*CF*_) compared to the previous two design iterations. We predicted that the self-circulating cross flow would prevent stacking of the waste intermediate-sized particles, allowing the small particles to follow a radial path through zone 3.

The AD zone from Fig. 2C was further simulated in COMSOL (Fig. 3A) to determine how different values of the angular displacement (1°, 3° and 5°) would affect the cross flow in the device. The cross flow outlet was left out from this simulation to solely focus on effect of the position of the pillars on cross flow velocity. The bottom panels in figure 3A show that the cross flow surface velocity (red zones) increased with increase in the displacement angle. The velocity was measured along the dotted green line in the inset of figure 3B. Fig. 3B shows a plot of the cross flow velocity as a function of the displacement angle for different flow rates (i.e. from 100 *µ*l/min to 500 *µ*l/min). For any given flow rate, the cross flow peaked at an angular displacement (*θ*) of 5°. This was expected as our pillar arrangement has a periodicity of 10^o^ (i.e. angular displacements of 0° and 10° are exactly the same). The final devices were fabricated with an angular displacement of 5°. For a given displacement angle, the cross flow also increased with an increase in the inlet flow rate.

**Figure 3:**
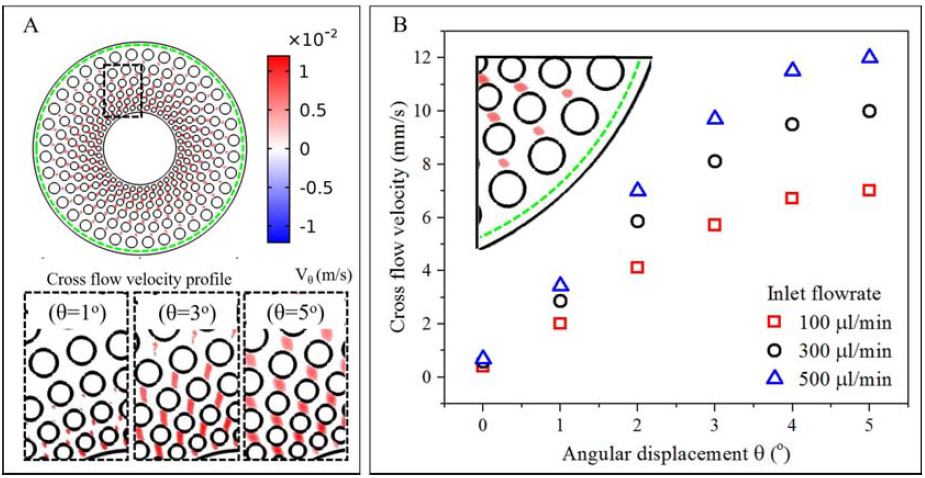
Results of COMSOL simulation to study the effect of the angular displacement on the magnitude of the cross flow velocity. (A) The fluid flow in the zone 2 (AD zone) was simulated for angular displacements (*θ*) of 1°, 3° and 5° respectively. The bottom panels show that the cross flow component (red areas) increases with an increase in the angular displacement. (B) The cross flow velocity was plotted against the angular displacement for different flow rates. The cross flow increases with increase in both angular displacement and the inlet flow rate. The velocity was measured along the dotted green line shown in the inset.

### 3.2 Clogging-free separation of both large and small particles in a single experiment

Figure 4 shows the paths of different particles in the device. As shown in panel (A), the 10 *µ*m beads were trapped by the zone 1 pillars encircling the inlet and they did not enter the device. The inlet region was 1.5 mm in diameter and I 5 mm (PDMS chip width) in height. Due to the large volume of the inlet region, the 10 *µ*m particles trapped at the inlet did not stop the device operation due to clogging. At the end of the experiment, buffer solution was flowed from the outlets to collect all the trapped 10 *µ*m beads from the inlet.

**Figure 4:**
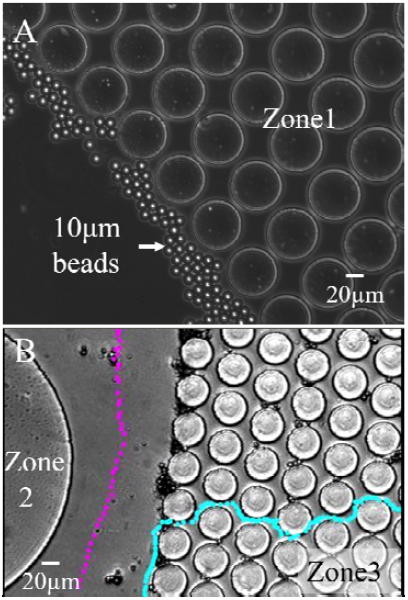
Paths taken by particles of different sizes in RAPID. (A) The 10 *µ*m beads are stopped at the inlet by the pill zone 1 and they do not enter the device. (B) The tracks followed by a 2 *µ*m (light-blue) and a 7 *µ*m (purple) partic shown. The small particle moves radially outward, while the large particle follows the cross flow.

As shown in the supplementary video SV1, initially the 7 *µ*m particles followed a radial path from zone 1 to zone 3. The 3 *µ*m pillar gap in zone 3 prevented the 7 *µ*m particles from going towards outlet O_*RF*_, and they started to collect before the first row of pillars in zone 3. The increased flow resistance in the radial direction further strengthened the cross flow between second and third zones. As shown in the video SV2, the strong cross flow carried away the remaining 7 *µ*m particles to the waste outlet. The 2 *µ*m beads took an approximately radial path through zone 3 towards the radial flow outlet. While no 7 *µ*m particle came out of the device through the radial path, some of the 2 *µ*m particles have remained in the cross flow. Panel (B) shows the paths taken by 7 *µ*m (purple) and 2 *µ*m beads (light-blue) after crossing zone 2. The experimentally obtained tracks of the particles agreed with the predictions from the COMSOL simulation of the fluid flow in the device.

The video SV3 confirms that the 2 *µ*m beads were able to make their way through zone 3 even in the presence of st 7 *µ*m beads. It is because the height of the device was 10 *µ*m, leaving the 2 *µ*m beads enough room to pass through the between the stacked 7 *µ*m beads and the pillars. It should be noted that the height of the device can be appropr adjusted depending on the size of the microparticles to be separated.

In summary, the self-sustaining cross flow is responsible for removing most of the intermediate-sized (7 *µ*m) beads RAPID, thereby allowing us to separate both large (10 *µ*m) and small (2 *µ*m) beads in a single experiment. The concent of the 10 *µ*m beads in this experiment was chosen to be 30000 times smaller than the other two kinds of particles experiment to simulate the capture of large rare cells. As discussed in the supplementary information, we also demons separation of concentrated small rare particles by mixing 7 *µ*m and 1 *µ*m beads in 10:1 ratio.

### 3.3 Purity, recovery and throughput

The performance of the particle sorting devices is typically measured in terms of throughput, purity and rec Throughput is defined by the total sample volume collected per unit time. Figure 5A shows that the throughput incr with increase in the inlet flow rate. The plot of throughput as a function of inlet flow rate confirms that RAPID is capa operating with almost no dead volume over a large range of flow rates. The plot also shows that the overall throughput filled square) is primarily dominated by that of the cross flow outlet (red open circle). This is because the cross flow pa a much lower hydrodynamic resistance than the radial path in our design.

**Figure 5:**
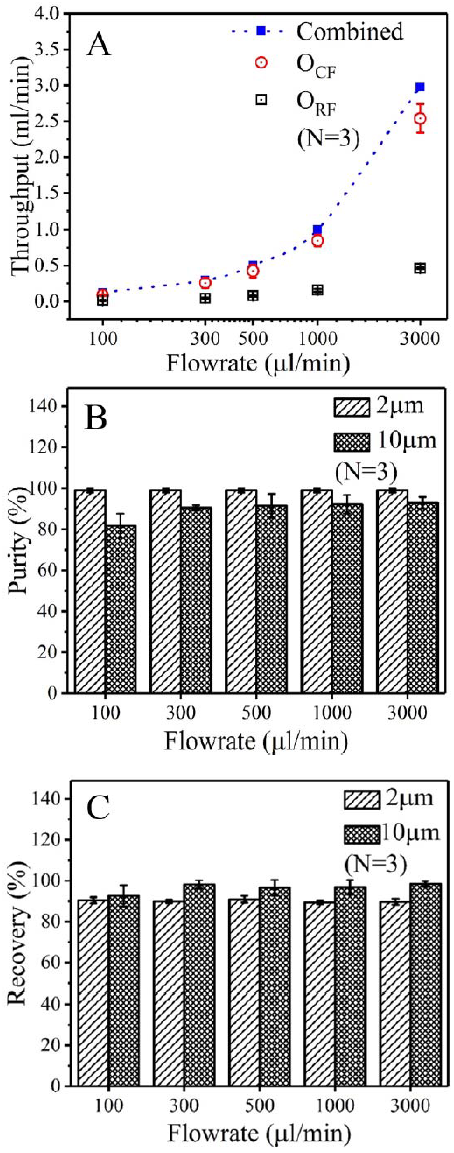
Separation pe*RF*ormance of small and large beads in RAPID. (A) The throughput of the device increases w increase in the inlet flow rate. The throughput of the radial flow outlet is always less than that of the cross flow because of the higher hydrodynamic resistance in the radial direction. (B) The purity of the 2 *µ*m beads at the radia outlet remains almost constant at 99.5% irrespective of the inlet flow rate. In contrast, as the inlet flow rate increase purity of the 10 *µ*m beads at the inlet increases from 80% to 95%. (C) The recovery of both small and large beads is a independent of the flow rate. The average recoveries of small and large beads are ⟂ 90% and ⟂ 96% respectively.

Hydrodynamic resistance in radial path is high, because it is determined by very small pillar gap in zone 3. We achie combined throughput (i.e. the sum of the throughput from individual outlets) of ⟂ 3 ml/min at the highest flow rat ml/min, thereby justifying the name RAPID.

Purity is the ratio of the number of the desired beads to the total number of beads at a particular inlet or outle instance, the 2 *µ*m beads are the desired beads at the radial outlet, whereas, the 10 *µ*m beads are the desired beads inlet. Figure 5 B shows the effect of the flow rate on the purity of these two kinds of beads. The radial flow outlet can maintain ⟂ 99.5% purity for the 2 *µ*m beads over a 30-fold increase in the inlet flow rate. This is because the 7 *µ*m beads cannot reach the radial flow outlet even at the highest flow rate. The purity of the 10 *µ*m beads at the inlet increases from ⟂ 80% to ⟂ 95% with an increase in flow rate from 100 *µ*l/min to 3000 *µ*l/min. The reason for the decreased purity of the 10 *µ*m beads at low flow rates is the occasional trapping of 7 *µ*m and 2 *µ*m beads behind 10*µ*m beads. An increase in the flow rate dislodges these stacked smaller beads, thereby improving the purity. We could not increase the flow rate beyond 3 ml/min because neither the connectors nor the PDMS-glass chip could withstand higher flows.

Recovery of a separation device indicates how many microparticles are successfully retrieved from the outlet. It is defined as the ratio of the total number of particles at the outlet and the total number of particles at the inlet. The number of each type of beads was counted before loading them in the syringe, and this number was taken as the number of beads in the inlet. We counted the total number of 2 *µ*m beads collected from both the outlets while calculating the recovery. The 10 *µ*m beads remaining in the inlet were collected using the reverse flow and this number was taken as the number of 10 *µ*m beads in the outlet. Figure 5C shows the recovery of 2 *µ*m and 10 *µ*m beads (N = 3) at different flow rates. As seen from the plots, the final recovery of the beads is independent of the flow rate. The average recoveries of the 2 *µ*m beads and 10 *µ*m beads are ⟂ 90% and ⟂ 96% respectively. As discussed in the supplementary section, the recovery rate of the particles is a strong function of the flow rate.

### 3.4 Comparison of RAPID with other pillar-based separation devices

Table 1 compares the pe*RF*ormance of RAPID with dead-end, cross flow and DLD pillar designs reported in the recent literature [22, 23, 24] for separation of both small (S) and large (L) beads. Both RAPID and DLD can achieve comparable high purity and recovery for both small and large particles simultaneously, unlike dead-end and cross flow devices. Finally, the throughput achieved in this RAPID prototype is 10 and 100 times higher than the crossflow and dead-end filter devices respectively, but is almost three times less compared to DLD device. The throughput of RAPID can be increased further by using high pressure connectors and a monolithic thermoplastic chip. RAPID can handle much higher samples concentrations compared to DLD [24]. The reason for this is that DLD relies on flow bifurcation. At high concentrations the particle to particle interaction increases which affect bifurcation and also leads to bead aggregation, this disturbs the flow profile and affects the DLD performance.

**Table 1:** Comparison of RAPID with other pillar-based filters for microparticle sorting

|  | Dead-end pillar[22] | Cross flow filter[23] | DLD [24] | RAPID |
| --- | --- | --- | --- | --- |
| Purity | 85-97% (S)<br>3-15% (L) | 97% (S)<br>80-97% (L) | 85-99% (S)<br>72-99%(L) | 99% (S)<br>81-91% (L) |
| Recovery | 48-80% (S)<br>20-52% (L) | 15-3% (S)<br>80-96% (L) | 96% (S)<br>99%(L) | 90% (S)<br>92-98% (L) |
| Throughput<br>(mL/min) | 0.02 | 0.1 | 10 | 3 |

S and L denote the corresponding parameters for small and large particles respectively.

## 4 Conclusions

Most of the dead-end pillar filters suffer heavily from clogging issues and tend to have extremely low throughput (tens to hundreds of *µ*l/min). On the other hand, traditional cross flow filter designs have high throughput and almost no clogging, but require a much larger footprint to increase device efficiency. Moreover, both small and large target cells continue to remain in the main channel along with other non-target cells. We designed a novel pillar-based filter with a radial geometry (RAPID) for simultaneous separation of small and large particles. It starts working like a dead-end pillar filter, but automatically turns into a cross flow device on the onset of clogging. It is the first size-based particle separation device to combine the advantages of both dead-end and cross flow device types to separate multi-sized particles. We tested continuous clog-free operation of the device up to 1 hour at the lower flow rates. We have achieved a throughput of 3 ml/min, which is 10 and 100 times higher than that reported by cross flow and dead-end filterrespectively, which justifies the acronym RAPID. We also achieved purity and recovery in excess of 90% for both small and large particles. Our pe*RF*ormance parameters are comparable to DLD devices, with an additional advantage of the ability to handle concentrated samples.

## acknowledgements

The authors would like to acknowledge the Centre for Nanoelectronics (phase 2) in IIT Bombay for partial funding and Dr. Dhrubaditya Mitra (NORDITA, Stockholm) for helpful discussions. Mr. Milan Khadiya for his support on counting. The devices have been fabricated in the cleanroom of the IIT Bombay Nanofabrication Facility.

## References

[1] C.C. Wu, L.Z. Hong, C.T. Ou, Journal of Medical and Biological Engineering 32(3), 163 (2012)

[2] X. Chen, C.C. Liu, H. Li, et al., Sensors and Actuators B: Chemical 130(1), 216 (2008)

[3] L.R. Huang, E.C. Cox, R.H. Austin, J.C. Sturm, Science 304(5673), 987 (2004)

[4] V. Murlidhar, M. Zeinali, S. Grabauskiene, M. Ghannad-Rezaie, M.S. Wicha, D.M. Simeone, N. Ramnath, R.M. Reddy, S. Nagrath, Small 10(23), 4895 (2014)

[5] S.M. MCFaul, B.K. Lin, H. Ma, Lab on a chip 12(13), 2369 (2012)

[6] A.A.S. Bhagat, H. Bow, H.W. Hou, S.J. Tan, J. Han, C.T. Lim, Medical & biological engineering & computing 48(10), 999 (2010)

[7] V. VanDelinder, A. Groisman, Analytical chemistry 78(11), 3765 (2006)

[8] T. Songjaroen, W. Dungchai, O. Chailapakul, C.S. Henry, W. Laiwattanapaisal, Lab on a Chip 12(18), 3392 (2012)

[9] A.J. Mach, D. Di Carlo, Biotechnology and bioengineering 107(2), 302 (2010)

[10] N. Pamme, Lab on a Chip 7(12), 1644 (2007)

[11] Y. Cheng, X. Ye, Z. Ma, S. Xie, W. Wang, Biomicrofluidics 10(1), 014118 (2016)

[12] S.B. Huang, M.H. Wu, G.B. Lee, Sensors and Actuators B: Chemical 142(1), 389 (2009)

[13] W. Liu, L. Li, J.c. Wang, Q. Tu, L. Ren, Y. Wang, J. Wang, Lab on a chip 12(9), 1702 (2012)

[14] S.B. Huang, Y. Zhao, D. Chen, H.C. Lee, Y. Luo, T.K. Chiu, J. Wang, J. Chen, M.H. Wu, Sensors and Actuators B: Chemical 190, 928 (2014)

[15] Y. Yoon, S. Kim, J. Lee, J. Choi, R.K. Kim, S.J. Lee, O. Sul, S.B. Lee, Scientific reports 6, 1 (2016)

[16] X. Chen, D. Cui, C. Liu, H. Li, J. Chen, Analytica chimica acta 584(2), 237 (2007)

[17] J. Happel, H. Brenner, Low Reynolds number hydrodynamics: with special applications to particulate media, vol. 1 (Springer Science & Business Media, 2012)

[18] E.D. Pratt, C. Huang, B.G. Hawkins, J.P. Gleghorn, B.J. Kirby, Chemical engineering science 66(7), 1508 (2011)

[19] D. Pappas, K. Wang, analytica chimica acta 601(1), 26 (2007)

[20] H.y. Li, C.D. Bertram, D.E. Wiley, AIChE journal 44(9), 1950 (1998)

[21] Z. Geng, Y. Ju, W. Wang, Z. Li, Sensors and Actuators B: Chemical 180, 122 (2013)

[22] J. Alvankarian, A. Bahadorimehr, B.Y. Majlis, Biomicrofluidics 7(1), 014102 (2013)

[23] Y.Y. Chiu, C.K. Huang, Y.W. Lu, Biomicrofluidics 10(1), 011906 (2016)

[24] D.W. Inglis, N. Herman, G. Vesey, Biomicrofluidics 4(2), 024109 (2010)

